# Predicting mechanisms of action at genetic loci associated with discordant effects on type 2 diabetes and abdominal fat accumulation

**DOI:** 10.1101/2022.04.27.489778

**Authors:** Yonathan Tamrat Aberra, Lijiang Ma, Johan L.M. Björkegren, Mete Civelek

## Abstract

Metabolic syndrome (MetSyn) is a cluster of dysregulated metabolic conditions that occur together to increase the risk for cardiometabolic disorders such as type 2 diabetes (T2D). One key condition associated with MetSyn, abdominal obesity, is measured by computing the ratio of waist-to-hip circumference adjusted for the body-mass index (WHRadjBMI). WHRadjBMI and T2D are complex traits with genetic and environmental components, which has enabled genome-wide association studies (GWAS) to identify hundreds of loci associated with both. Statistical genetics analyses of these GWAS have predicted that WHRadjBMI is a strong causal risk factor of T2D and that these traits share genetic architecture at many loci. To date, no variants have been described that are simultaneously associated with protection from T2D but with increased abdominal obesity. Here, we used colocalization analysis to identify genetic variants with a shared association for T2D and abdominal obesity. This analysis revealed the presence of five loci associated with discordant effects on T2D and abdominal obesity. The alleles of the lead genetic variants in these loci that were protective against T2D were also associated with increased abdominal obesity. We further used publicly available expression, epigenomic, and genetic regulatory data to predict the effector genes (eGenes) and functional tissues at the 2p21, 5q21.1, and 19q13.11 loci. We also computed the correlation between the subcutaneous adipose tissue (SAT) expression of predicted effector genes (eGenes) with metabolic phenotypes and adipogenesis. We proposed a model to resolve the discordant effects at the 5q21.1 locus. We find that eGenes gypsy retrotransposon integrase 1 (*GIN1*), diphosphoinositol pentakisphosphate kinase 2 (PPIP5K2), and peptidylglycine alpha-amidating monooxygenase (*PAM*) represent the likely causal eGenes at the 5q21.1 locus. Taken together, these results are the first to describe a potential mechanism through which a genetic variant can confer increased abdominal obesity but protection from T2D risk. Understanding precisely how and which genetic variants confer increased risk for MetSyn will develop the basic science needed to design novel therapeutics for metabolic syndrome.

## INTRODUCTION

Metabolic syndrome (MetSyn) is a cluster of dysregulated metabolic conditions that tend to occur together to increase the risk for cardiometabolic disorders such as type 2 diabetes (T2D)^1^. This cluster includes insulin resistance (IR), abdominal obesity, elevated serum triglycerides (TG) levels, low high-density lipoprotein cholesterol (HDL-C) levels, as well as elevated systolic and diastolic blood pressure. Obesity, or the excessive accumulation of fat that presents a risk to health, is a major contributor to MetSyn^1,2^. Obesity, which is typically defined as a Body-Mass Index (BMI) above 30, has reached unprecedented levels of prevalence, and its role as a central regulator of disease risk makes it an appealing therapeutic target^2^. Several recently developed T2D therapeutics have even successfully targeted obesity; SGLT2 inhibitors and GLP-1 agonists have been reported to result in a 2-6 kilogram reduction of body weight and reduced insulin resistance^3^.

Despite the promise of these obesity-centered therapeutic strategies, there has also been a growing body of evidence describing a rare phenotype known as metabolically healthy obesity (MHO)^4^. MHO describes a group of phenotypes in which individuals with obesity are protected from adverse metabolic effects^5^. While no formal definition of MHO exists, it is often described as either obesity with less than three components of MetSyn, or obesity without insulin resistance as computed by the Homeostasis Model Assessment of Insulin Resistance (HOMA–IR)^5^. Mechanisms proposed to mediate this include depressed ectopic fat accumulation, subcutaneous adipose tissue expansion plasticity, and shifts in fat storage from the abdomen to the legs^4–6^. In recent years the ability of abdominal obesity to mediate cardiometabolic disease risk has gained attention. People with MHO have less intra-abdominal fat accumulation compared to people with metabolically unhealthy obesity (MUO)^7–12^. Intra-abdominal fat accumulation can be approximated through the ratio of Waist-to-Hip circumference (WHR) adjusted for BMI (WHRadjBMI). WHRadjBMI is a causal factor that increases susceptibility for T2D, but the genetic and molecular mechanisms underlying fat distribution remain largely unknown^13–15^. Understanding the mechanisms mediating WHRadjBMI, MHO, and T2D is critical to our understanding of disease pathogenesis and to clinical strategies to treat MetSyn.

Most of the genetic mechanisms of MHO described have been associated with increased BMI without increased disease risk. For example, the missense variant rs373863828 in CREB3 Regulatory Factor has been shown to increase BMI without a corresponding increase in HOMA-IR and circulating triglycerides, or a decrease in circulating adiponectin^16^. Ob/ob mice with overexpression of adiponectin but lacking in leptin are shown to accumulate considerable fat mass without a corresponding increase in insulin sensitivity^17^. In contrast, the genetic loci associated with increased WHRadjBMI but without increased disease risk have not yet been described. To date, all genes that have been shown to increase fat accumulation into abdominal fat depots have also been shown to increase the risk for T2D^18–23^.

As complex traits with both environmental and genetic risk factors, abdominal obesity and T2D have been the subject of multiple genome-wide association studies (GWAS). While GWA studies have identified hundreds of genetic loci associated with abdominal obesity and T2D, moving from association to mechanism at a locus is not trivial. The use of colocalization analysis (COLOC), which identifies loci that contain shared genetic architecture for multiple traits of interest, can inform mechanistic hypotheses moving from association to function by integrating data from multiple studies^24–27^. For example, the colocalization of a GWAS signal with genetic regulation of genes at quantitative trait loci (QTL) implies a mechanistic relationship between the regulated gene and GWAS trait^28^. Another recently developed approach named Tissue of ACTion scores for Investigating Complex trait-Associated Loci (TACTICAL)^29^, incorporates gene expression data, and epigenetic annotations with GWAS associations to predict the causal eGenes and tissues of action at GWAS loci. These methods have been used to inform data-driven mechanistic predictions at GWAS loci that have been experimentally validated and can recall previously validated loci as positive controls.

To advance the understanding of mechanisms linking body fat distribution to T2D risk, independent of overall obesity, we used COLOC and TACTICAL to predict the mechanisms of action at genetic loci associated with both T2D and WHR, both adjusted for the BMI. Using the most recent GWAS summary statistics, QTL summary statistics, tissue-specific gene expression data, and high-resolution epigenetic annotations, we predicted the shared genetic architecture of T2DadjBMI and WHRadjBMI at 79 genetic loci. Here we present the identification of 5 loci that contained association signals with discordant effects on abdominal fat and T2D risk, meaning that the allele of the lead variant associated with protection from T2D was associated with increased abdominal fat accumulation. We predicted the eGenes and tissues of action at these 5 loci and explored the relationship between adipose eGene expression with cellular and physiological phenotypes. Here, we provide data-driven hypotheses about predicted candidate causal eGenes at GWAS loci with associations that recall metabolically healthy abdominal obesity.

## RESULTS

### Colocalization analysis of genetic loci associated with Type 2 Diabetes and Body Fat Distribution predicts colocalization of discordant T2DadjBMI and WHRadjBMI association signals at six loci

To identify genetic loci which contained pleiotropic association signals for both T2DadjBMI and WHRadjBMI, we performed colocalization analysis (Figure 1A). This analysis yielded 79 genetic loci where a single variant was significantly associated with both T2DadjBMI and WHRadjBMI. We obtained the 99% credible set of variants in colocalized loci (Supplementary File 1) and discovered the presence of 143 variants in five loci associated with discordant effects on T2DadjBMI and WHRadjBMI. We also discovered 851 SNPs in 73 loci with the expected concordant effects on both traits. Although almost all of the representative lead discordant variants reached genome-wide significance, two associations reached nominal significance (p < 5e-05). Because of recent work demonstrating that even variants with only nominal and local significance in GWAS can also have functional relevance to GWAS traits, we included variants prioritized in the 99% credible set but with only nominal significance^30^. We then performed fine-mapping of the causal variants in each locus containing a discordant association signal while relaxing the assumption of a single causal variant per locus. In four of the five loci, this fine-mapping recalled only one likely candidate causal signal. In the 5q21.1 locus, SuSiE identified a secondary association signal that was also associated with discordant effects on T2DadjBMI and WHRadjBMI (Figure 1B and Supplementary File 2). To parse the associations between specific components of WHRadjBMI, including WC, HC, WHR, and BMI, with both T2D and T2DadjBMI, we performed multi-trait colocalization analysis with Hyprcoloc of the associations at discordant loci (Supplementary File 3). At three of the five discordant loci, the discordant association signals were also colocalized with WHRadjBMI component traits waist circumference and WHR.

**Figure 1.**
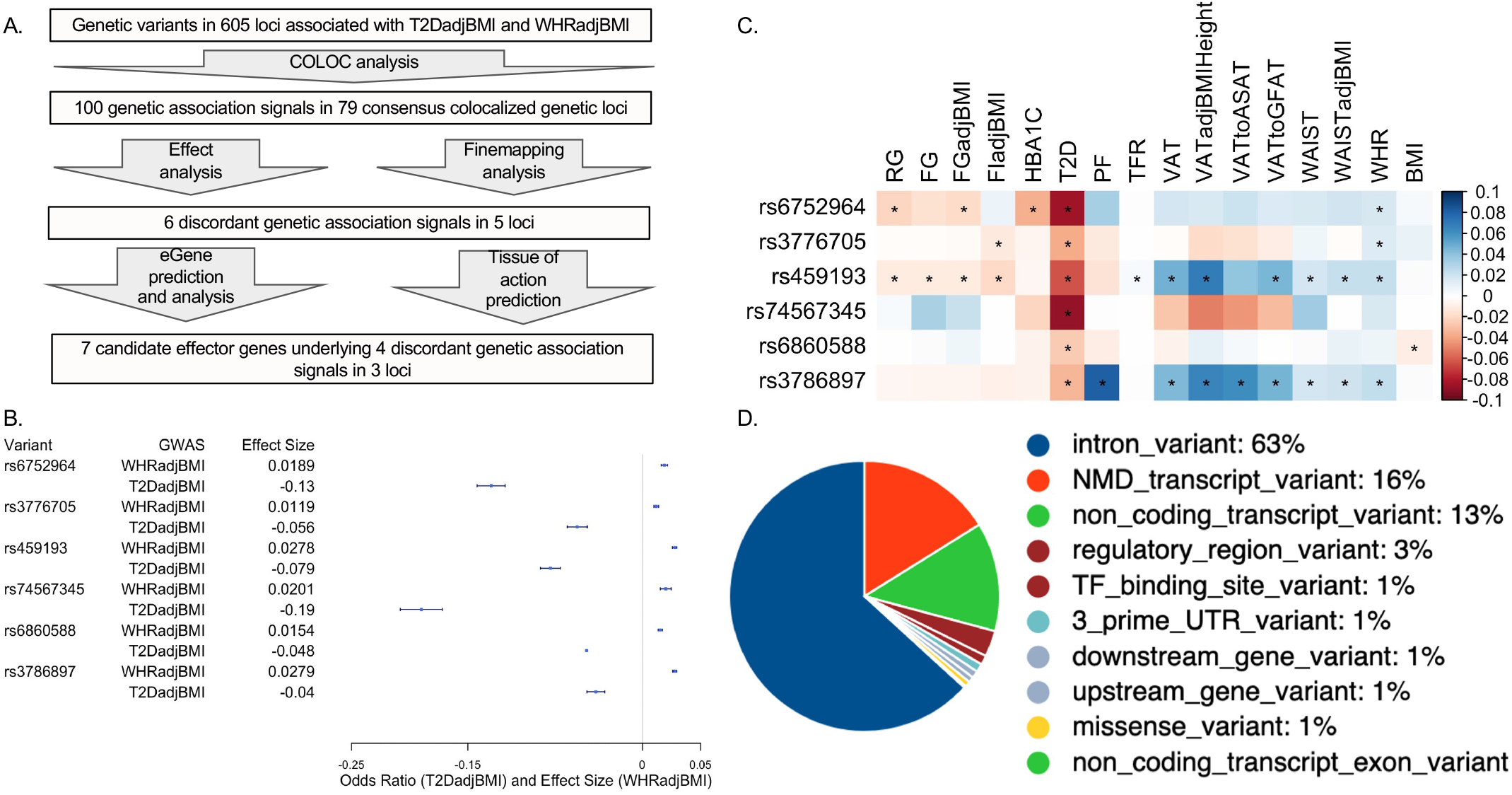
Analysis summary and discordant variant characteristics. (A) Summary of analysis pipeline and generated results. Details of data sources are available in Supplementary Table 1. (B) Effect size (WHRadjBMI) and Odds Ratio (T2DadjBMI) of lead genetic variant at discordant association signals. (C) PheWAS of lead discordant genetic variant effect sizes on glycemic and anthropometric traits. From left to right: Random Glucose (RG), Fasting Glucose (FG), FG adjusted for BMI (FGadjBMI), Fasting Insulin adjusted for BMI (FIadjBMI), Glycated Hemoglobin (HBA1C), Pancreatic Fat percentage (PF), Trunk Fat Ratio (TFR), Visceral Adipose Tissue (VAT), VAT adjusted for BMI and Height (VATadjBMIHeight), VAT to Abdominal Subcutaneous Adipose Tissue (VATtoASAT), VAT to Gluteofemoral Fat (VATtoFGAT), Waist circumference, Waist Circumference adjusted for BMI (WCadjBMI), Waist-to-Hip Ratio (WHR), and BMI. (D) Variant effect prediction of 99% credible set variants in discordant genetic loci.

We next investigated the genetic and physiological consequences of discordant variants. We performed a phenome-wide association study (PheWAS) for anthropometric and glycemic traits with the most highly powered GWAS available (Figure 1 – source data 1)^31^. We used the most highly powered GWAS or GWAS meta-analysis for each trait included in our PheWAS and queried the summary statistics for the associations of each lead discordant variant (Figure 1C). This query revealed consistent significant associations with discordance across anthropometric and glycemic traits in each locus. At the association signal in the 5q11.2 region, association signals exemplified this metabolic discordance. Represented by genetic variant rs459193, the association signal was associated with increased abdominal obesity in nearly every metric, but also with protection from type 2 diabetes in nearly every metric. At all lead discordant variants, effects were consistent with a phenotype of increased abdominal obesity but protection from type 2 diabetes.

We then queried the Variant effect predictor (VEP) to discover genetic variant annotations^32^ (Figure 1D and Supplementary File 4). VEP predicted that discordant variants overwhelmingly lie in noncoding regions of the genome, with only one missense variant in a coding region. Because the vast majority of discordant variants lie in noncoding regions, it is likely their function lies in altering genetic regulation of proximal genes^33^. Therefore, we investigated the coincidence of these discordant variants with the genetic regulation of proximal genes with functional prediction methods.

### Integration of molecular QTLs and genomic annotations to predict functional genes in tissues of action at discordant genetic loci

To investigate the role of eGenes in physiological phenotypes and cellular phenotypes, we evaluated the correlation of adipose tissue eGene expression and T2D-relevant phenotypes since these correlations can reveal biologically-relevant functional relationships^34^. To predict the genes and tissues of function at discordant loci, we used publicly available multi-omic data from metabolically relevant tissue-specific resources to predict functional mechanisms underlying associations. We first interrogated where the 143 discordant variants in the credible set were located in relation to tissue-specific chromatin state data in pancreatic islet, adipose, liver, and skeletal muscle tissues^17^. We computed the enrichment of colocalized association signals in various chromatin state annotations in each of these tissues (Figure 2A). We noted the specific enrichment of adipose tissue chromatin states of high activity, such as active transcription start sites, enhancer regions, and areas of transcriptional activity. For every other tissue, the leading annotations represented areas of decreased transcriptional activity. We additionally queried 3D chromatin data for discordant variant enhancer/promoter contact but did not find any significant interactions (Figure 2 – source data 2). We then used these enrichment scores, chromatin states, and gene expression data to predict the functional tissues at each colocalized locus (Supplementary File 5). We predicted that adipose tissue was classified as the candidate TOA at three loci, and skeletal muscle and liver tissue shared classification with adipose tissue at the remaining two discordant loci (Figure 2B).

**Figure 2.**
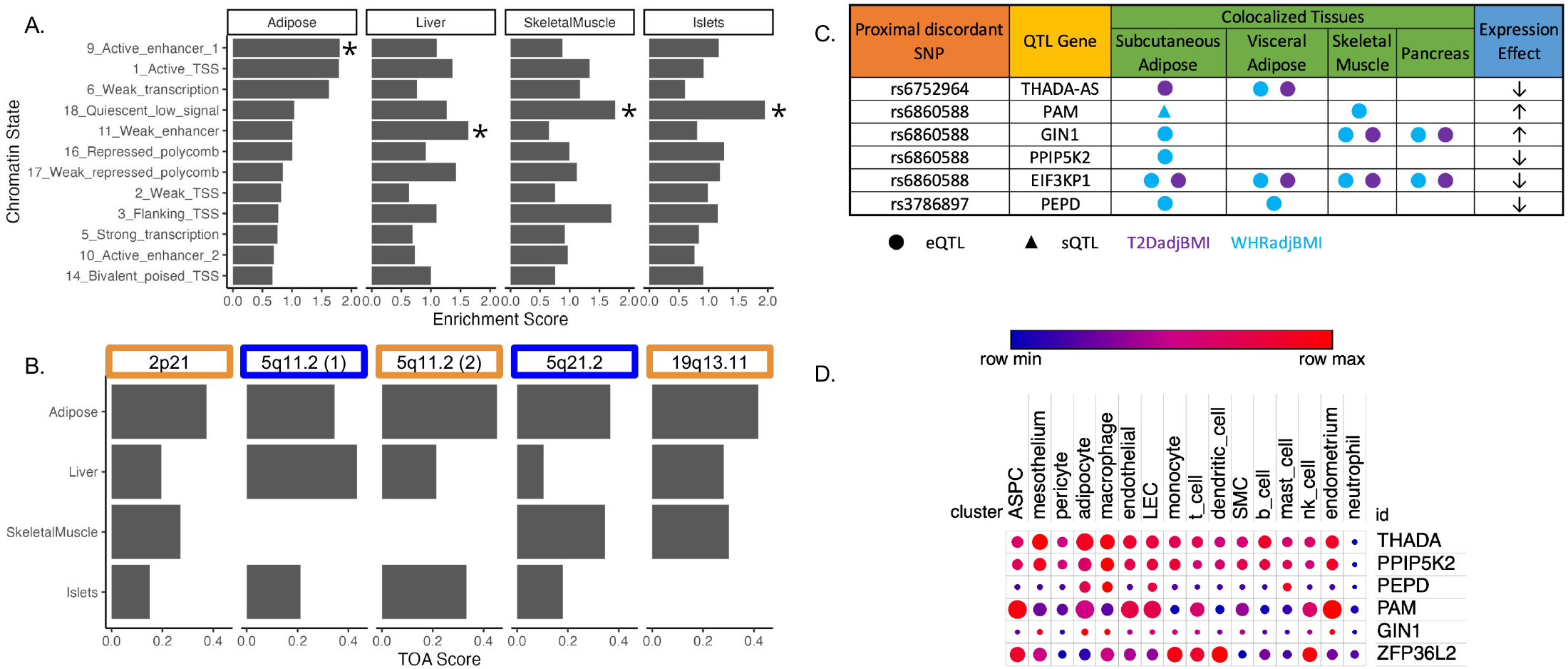
Predicting functional tissues and effector genes at discordant loci. (A) Tissue-specific enrichment of chromatin-states of variants in the 99% credible set of colocalized variants. (B) Tissue of Action scores for association signals in the six discordant loci. Orange coloration indicates predicted adipose tissue of action at the locus, and blue coloration indicates shared tissue of action assignment at the locus. (C) Summary table of the eQTL and sQTL colocalizations with WHRadjBMI and T2DadjBMI for discordant loci. The expression effect direction is with respect to the protective type 2 diabetes allele. (D) Expression of predicted effector genes in discordant loci across cell types. From left to right: Adipocyte progenitor stem cell (ASPC), Lymphatic endothelial cells (LEC), Smooth muscle cell (SMC). Data was obtained from Emont et al^35^.

To predict effector genes (eGenes) regulated by discordant variants, we next predicted the colocalization of quantitative trait loci (QTL) with the WHRadjBMI and T2DadjBMI GWAS. Colocalization of a GWAS association signal with a genetic regulatory association signal can be used to prioritize mechanisms underlying association. We obtained expression QTL (eQTL) and splicing QTL (sQTL) summary statistics from multiple cohorts and tissue groups (Figure 2 – source data 2). We extracted eQTL summary statistics for all genes within 1Mb of the lead variant of all discordant colocalized loci from adipose, pancreatic, skeletal muscle, and liver tissues. We extracted sQTL summary statistics for all genes within 1Mb of the lead variant of all discordant colocalized loci for adipose tissue data that was available. We used Summary-based Mendelian Randomization and Coloc.abf to perform GWAS-QTL colocalization and used the framework developed by Hukku et al. to reconcile the results of SMR and Coloc.abf. In this framework, colocalization found using Coloc.abf but not with SMR potentially represents signals with horizontal pleiotropy, whereas colocalization found through SMR but not through Coloc.abf potentially represents locus-level colocalization^24^. Colocalization found using both methods represents the identification of candidate causal effector transcripts. Our colocalization analysis revealed seven candidate causal effector transcripts at three of the 5 discordant loci (Figure 2C). With Coloc.abf, we predicted four putative eGenes in these two loci. At the 2p21 locus, we predicted *THADA-AS* (SAT, VAT) to be the sole eGene. At the 5q21.1 locus, we predicted *GIN1* (SAT), *PAM*(SAT & SKM), and *PPIP5K2* (SAT) to be the eGenes. The association signal at rs6860588 was associated with a novel alternative splicing isoform of *PAM* in subcutaneous adipose tissue (SAT), which skips the 14^th^ exon. Using SMR, we predicted four eGenes at two discordant loci. At the 5q21.1 locus, we predicted the genetic association signal represented by rs6860588 was also associated with the regulation of *EIF3KP1* (SAT, VAT, SKM, PANC), *PPIP5K2* (SAT), and *GIN1* (SAT, SKM, PANC). At the discordant association signal in the 19q13.11 locus, we predicted that the genetic association signal represented by variant rs3786897 was also associated with the regulation of *PEPD* (SAT, VAT). As the colocalization transcripts *GIN1* and *PPIP5K2* were replicated with both methods (Supplementary Files 6 and 7), these represent high-confidence predictions of potentially causal effector transcripts underlying the genetic association with discordance in the 5q21.1 locus. We queried white adipose tissue single-cell RNA sequencing data^35^ for discordant association signal eGenes and found that eGenes were expressed in adipocytes and adipocyte progenitor stem cells (ASPCs) (Figure 2D). Because body fat distribution associations are driven by ASPCs and adipocytes in adipose tissues^36–38^, we reasoned that exploring adipose expression data could help to explain discordant associations. This multi-omic data enabled us to make high-confidence consensus predictions of tissues and eGenes of action at discordant loci.

### Adipose gene expression analysis of discordant loci eGenes reveals dynamic expression in adipogenesis and relationships with metabolic physiology

To investigate the role of eGenes in physiological phenotypes and cellular phenotypes, we then evaluated the gene expression dynamics of eGenes in adipose tissue. Correlations between relevant tissue gene expression and metabolic phenotypes can reveal biologically-relevant functional relationships^34^. We used SAT transcriptomic data from the 426 men of the METSIM cohort to investigate how adipose tissue expression of discordant locus eGenes was related to 23 metabolic phenotypes underlying T2D and abdominal fat accumulation (Figure 3 – source data 3)^39,40^. We extracted adipose tissue gene-expression data for eGenes. Gene expression data were available for six of the seven eGenes. We additionally extracted splice junction expression data for the only gene with a colocalized splice junction, *PAM*. We then computed the biweight midcorrelation of transcript counts or splice junction counts with 23 metabolic phenotypes. We found significant (FDR < 0.05) correlations of adipose tissue gene expression of three genes with thirteen phenotypes (Figure 3A). We found that adipose tissue expression of *THADA-AS, PEPD*, and *GIN1* was significantly correlated with inflammatory, glycemic, and anthropometric phenotypes. SAT *THADA-AS* expression was positively correlated with insulin resistance, abdominal fat accumulation, and serum triglyceride levels, but with higher levels of plasma Interleukin-1 receptor antagonist (IL-1RA) and C-reactive protein (CRP). IL-1Ra plays a protective role in resolving inflammation^41^, and elevated levels have been linked to prediabetes^42,43^. CRP has been used as a biomarker of increased inflammation in chronic diseases^44^. The eQTL and GWAS data are associated with decreased expression of THADA-AS, which is consistent with the protection from insulin resistance in the correlation data but not with the increased abdominal obesity and inflammation. We are unable to resolve this correlation evidence with the discordance, but because the METSIM cohort was collected using single-end RNA sequencing, parsing the correlations of THADA and THADA-AS is difficult^45^. SAT expression of *GIN1* was correlated with higher plasma adiponectin. Adiponectin, secreted by adipocytes, increases insulin sensitivity, and this provides a mechanism for protection from T2D^46^. This expression is consistent with the QTL and GWAS data, providing a direct potential mechanism linking the eQTL to protection from T2D. SAT *PEPD* expression was also positively correlated with plasma IL-1RA levels. The QTL at this locus is associated with decreased expression of PEPD, providing another direct potential mechanism linking the eQTL to protection from T2D. Through this correlation analysis, we were able to predict the physiological consequences of eGenes at three discordant loci.

**Figure 3.**
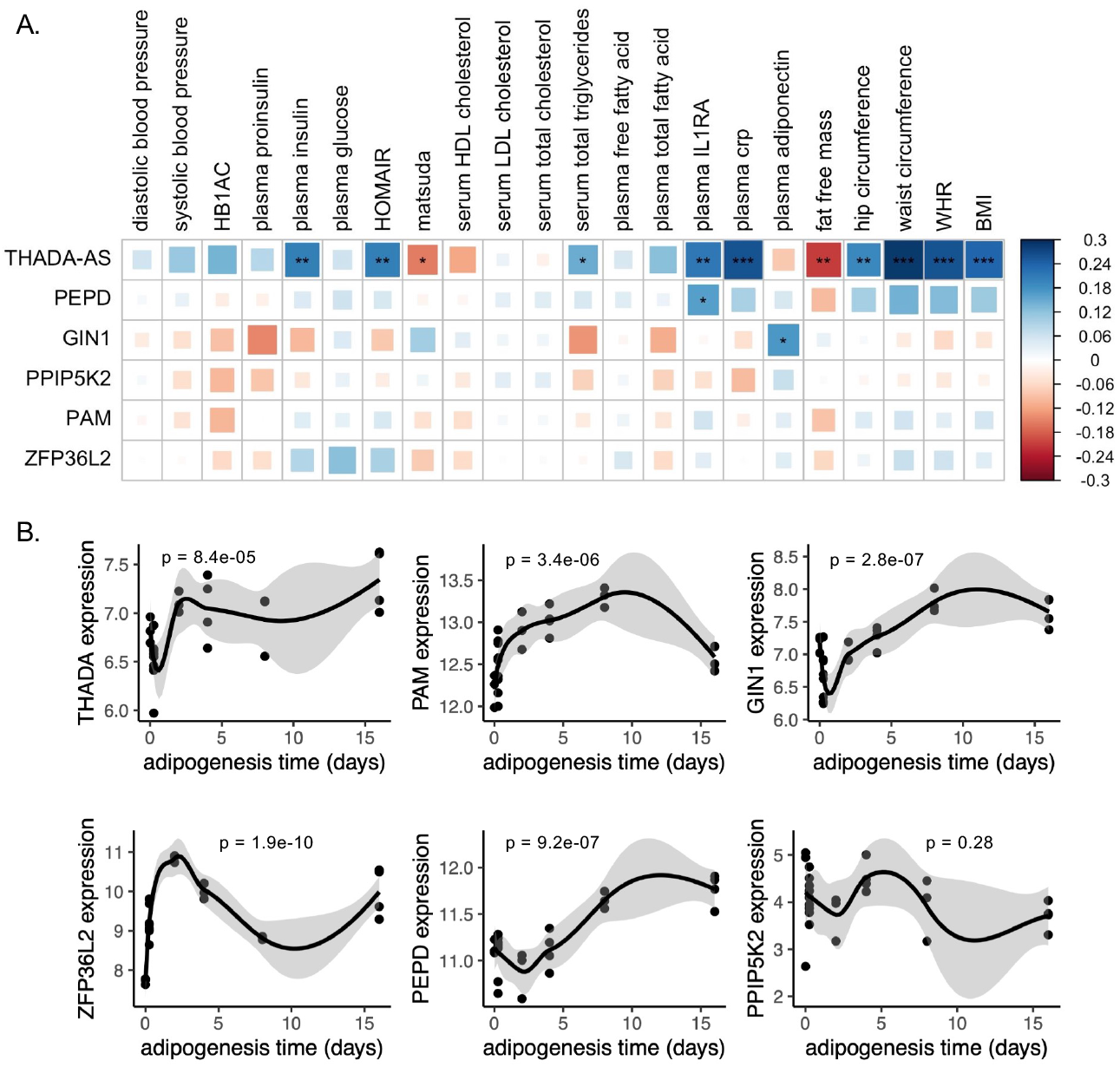
Predicted physiological and cellular effects of eGenes on metabolic phenotypes and adipogenesis. (A) Biweight midcorrelation of adipose tissue eGenes expression with metabolic phenotypes (FDR < 5%). From left to right: Homeostatic model of insulin resistance (HOMAIR), High-density lipoproteins (HDL), Low-density lipoproteins (LDL), Interleukin-1 Receptor Agonist (IL1RA), C-reactive protein (crp). (B) Dynamic expression of adipose tissue eGenes over sixteen-day adipogenesis time-course in Simpson-Golabi-Behmel syndrome (SGBS) cells. We performed the likelihood ratio test (LRT) to evaluate if each gene was dynamically expressed over the time course. The p-value of the LRT is included.

We next evaluated if eGenes identified in adipose tissues were dynamically expressed in adipogenesis. Dynamic gene expression in adipogenesis could point to the regulatory and structural roles of eGenes in adipogenesis^47,48^. We obtained time series ASPC adipogenesis time course data and evaluated eGenes for dynamic expression. Gene expression data were available for five of the seven eGenes. Because the expression data was single-stranded and unable to resolve forward or reverse-strand sequences, we included the probe for *THADA* to represent *THADA-AS*. We found that all eGenes except PPIP5K2 were dynamically expressed over a sixteen-day adipogenesis time course (Figure 3B), implying potential functional roles for these genes in regulating preadipocyte fate.

### Integration of analysis to predict the functional genes and tissues of action at the discordant 5q21.1 locus

By predicting the mechanisms of action at discordant loci, we were able to generate specific hypotheses about the genes at each locus that underlie GWAS associations. We predicted that the causal discordant signal at the 5q21.1 locus was represented by variant rs6860588. The T allele of rs6860588 is associated with protection from type 2 diabetes, increased abdominal obesity, decreased SAT expression of *GIN1*, increased SKM expression of *PAM*, decreased SAT expression of *PPIP5K2*, increased SAT expression of a *PAM* splice variant with a skipped exon 14, and decreased SAT expression of the canonical *PAM* splice junctions, exon 13:14 and exon 14:15 (Figure 4A, Figure 4-figure supplement 1, and Figure 4 - source data 4). While the eGenes, *PAM, GIN1*, and *PPIP5K2*, have not been studied in the context of obesity and metabolism, they have been studied for their function in other cell types. We found that GIN1 and PAM were dynamically expressed over the course of adipogenesis (Figure 3B). *GIN1* has been hypothesized to be a key regulator of energy metabolism in atria^49^, but little is known about gypsy integrases and their molecular function. PAM facilitates C-terminus glycine residue amidation, which can catalyze protein potency^50,51^. PAM additionally has been linked to metabolic phenotypes in multiple model organisms, where its deficiency is associated with decreased peptide secretion and potency critical to insulin release, but not with increased diabetes^52,53^. PAM loss of function likely results in deficient peptide synthesis and secretion in adipocytes as well, and its increase of function likely results in increased myokine signaling from skeletal muscle. Knockdown of PPIP5Ks results in decreased proliferation, increased mitochondrial mass, decreased inositol metabolism, and accelerated glycolysis in tumor cell lines^54–56^. We did not observe significant interactions between adipose *PPIP5K2* expression and adipogenesis or metabolic phenotypes, but this does not rule out a role for PPIP5K2 in the metabolic discordance at 5q21.1. Thus, we propose that the T allele at rs6860588 regulates a group of genes that promotes adipogenesis, glycolysis, and inflammation in white adipose tissue while simultaneously decreasing preadipocyte expansion and increasing skeletal muscle peptide secretion and potency (Figure 4B). This model is consistent with the tissue of action score and QTL analysis, which both predict skeletal muscle and adipose tissue contribution to the associations at the locus and reconcile the associations with abdominal obesity but protection from type 2 diabetes associated with the T allele of rs6860588 (Figure 4C).

**Figure 4.**
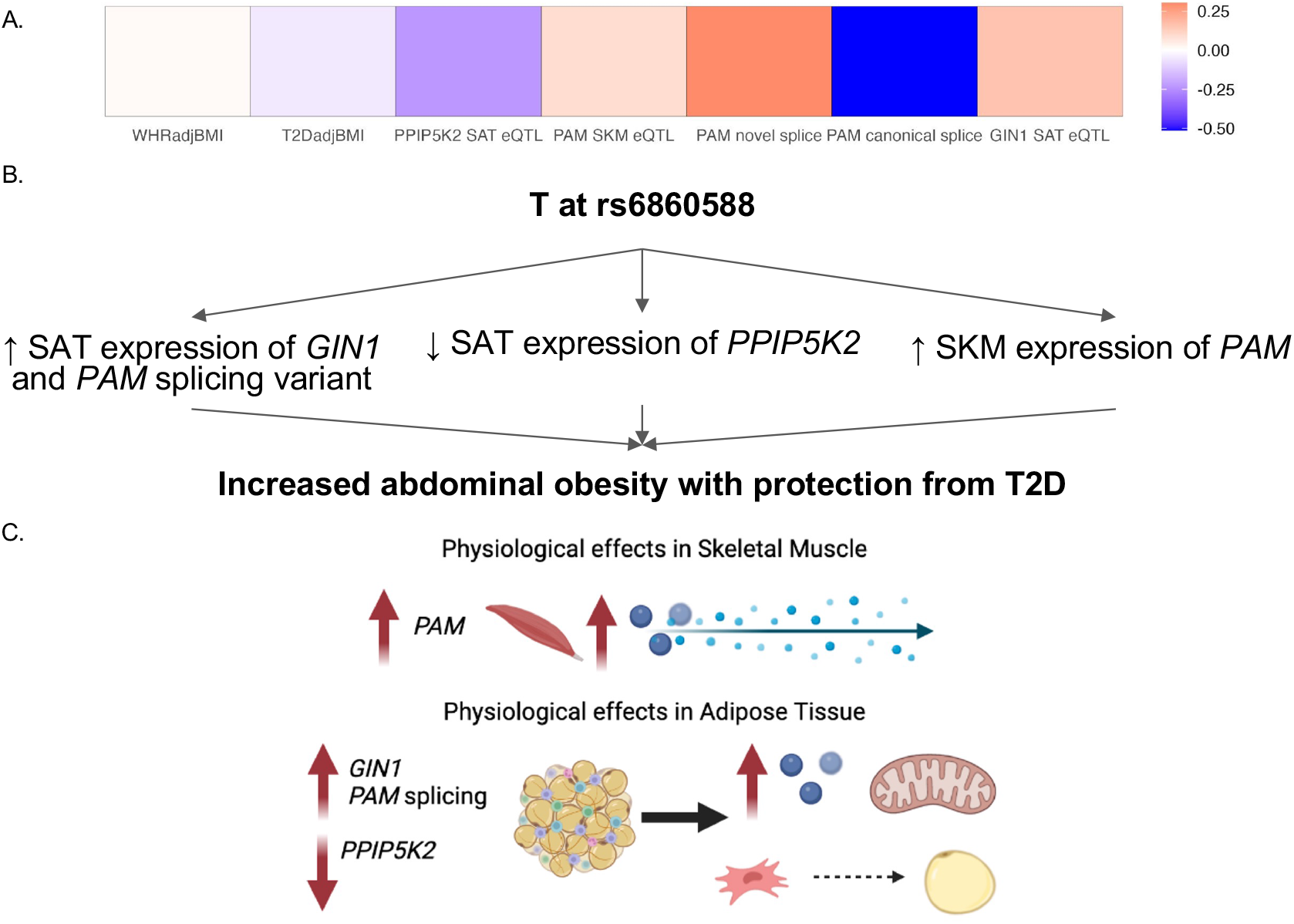
Predicted model of effects associated with T allele at rs6860588. (A) β of the T allele of discordant variant rs6860588 with respect to WHRadjBMI, T2DadjBMI, and colocalized eGenes. (B) Summary of associations with T allele at rs6860588. (C) Integrated model reconciling metabolic discordance with eGene-associated phenotypes in two tissues of action. Created with BioRender.com.

## DISCUSSION

We report here the integration of multi-omic data spanning the genome, transcriptome, and epigenome to predict functional genes and tissues underlying genetic signals associated with abdominal obesity but protection from T2D. We predicted the colocalization of T2DadjBMI and WHRadjBMI association signals at 79 genetic loci. The protective allele of six association signals was associated with lower T2D risk but higher abdominal fat accumulation, independent of overall obesity (Figure 1). By analyzing colocalization with molecular QTLs, computing the enrichment of variants in epigenomic and genomic annotations, and comparing tissue-specific gene expression, we predicted the eGenes and tissues of action at discordant association signals (Figure 2). We found significant evidence that adipose tissue biology is a significant contributor at colocalized loci. We then explored the effects of eGenes expression in adipose tissue and preadipocytes on adipogenesis metabolic phenotypes (Figure 3) and proposed a model by which the genetic variant rs6860588 might confer protection from T2D but increased abdominal obesity (Figure 4).

The six genetic association signals associated with discordant metabolic phenotypes offer potential insight into the genetic mechanisms underlying risk stratification of T2D risk within abdominal obesity. While mechanisms promoting MHO have been described, most have focused on body fat distribution. Defining more mechanisms that promote MHO is critical as rates of obesity rise globally. Complicating the study of MHO is the lack of precision in its definition. Some definitions include obesity with less than three components of MetSyn, obesity with healthy HOMA-IR, or even obesity with the lack of a metabolic and cardiovascular disorder^4^. MHO has been controversial and termed an intermediate state^57–59^, but a growing body of evidence has accumulated providing evidence that genetic mechanisms influence predisposition to it. In Samoans, the common *CREBRF* coding variant rs12513649 increases BMI and overall adiposity but protects from insulin resistance^30^. Additionally, *IRS1, COBLL1, PLA2G6*, and *TOMM40* have been associated with higher BMI but with protective lipidemic and glycemic traits^6^. The physiological functions of these genes have been proposed to involve adipose tissue caloric load capacity and body fat distribution^6,36,60,61^.

While abdominal fat accumulation is known to be one of the strongest predictors of obesity-related complications^13,62,63^, our findings point to mechanisms that contradict this trend. Each locus must be functionally annotated before translating the association results to the clinic. If these discordant variants are functionally annotated and fully characterized, they might have clinical utility to T2D risk allele carriers and inform personalized therapeutic strategies. Discovering mechanisms uncoupling abdominal obesity from T2D can aid in personalized therapeutic strategies and in understanding personalized risk stratification. Risk-stratified personalized obesity treatment could prioritize patients that would or would not benefit significantly from weight-loss interventions, and use genotype as a biomarker for patients who would benefit from other therapeutic strategies^64,65^. Thus, the importance of personalized risk stratification for T2D will only increase as abdominal obesity becomes more prevalent. Personalized risk stratification with an understanding of specific molecular, cellular, and physiological mechanisms will aid in the prioritization of effective therapies. This investigation provided specific hypotheses linking functional genes at discordant loci to tissues of action for experimental follow-up *in vitro* and *in vivo*. Functional characterization of the effect of these genes on insulin uptake, preadipocyte proliferation, and adipogenesis, as well as secretome characterization, will elucidate precise mechanisms through which these eGenes might contribute to the discordant association signals.

We predicted tissues and mechanisms of action at five loci containing six discordant association signals with increased abdominal obesity and protection for type 2 diabetes. A particular example of a peculiar metabolic discordance was revealed at the 2p21 locus containing *THADA* and *THADA-AS*, represented by variant rs6752964 (Figure 4-figure supplement 2). The associations have been replicated multiple times^66–68^, but the exact mechanisms underlying this association are unknown. *THADA* plays an evolutionarily conserved role in intracellular calcium signaling and consequently non-shivering thermogenesis. In Drosophila melanogaster, *thada* knockout flies developed obesity and hyperphagia without altered circulating glucose levels^69^. In mice, pancreatic *Thada* knockout resulted in protection from T2D through the preservation of β-cell mass and improvement of β-cell function^70^. Mendelian randomization studies in humans have likewise found consistent links between THADA and adiposity, but have not yet been able to link it to diabetic phenotypes such as insulin secretion^68,71^. Our investigation revealed relationships between THADA and THADA-AS expression with diabetic and obesity-abdominal obesity phenotypes as well as dynamic expression in adipogenesis (Figure 3). Regulatory interactions whereby *THADA-AS* expression interferes with *THADA* transcription could provide a basis by which variant rs6752964 might confer abdominal obesity, but protection from type 2 diabetes^72–74^. Further, we also found colocalization of genetic regulation of *PEPD* in adipose tissue with the discordant association signal represented by variant rs3786897. Depletion of *PEPD* in preadipocytes has been shown to reduce adipogenic potential, decrease triglyceride accumulation, and phospho-Akt signaling, which is critical to insulin sensitivity^75^. Notably, a secondary signal represented by variant rs731839 was apparent in this locus but was not significant for WHRadjBMI. This signal has been associated with sex-specific effects on serum lipid levels in Han and Mulao populations^76^. Further *in vivo* and *in vitro* work must be done to resolve this multi-tissue, multi-eGene locus.

Although our analysis incorporated genome, transcriptome, epigenome, and phenome data in multiple cohorts, and used the consensus of orthogonal methods to predict the mechanisms of action at discordant loci, follow-up is required to validate each prediction. Additionally, our genetic expression data used single-strand sequencing, and therefore parsing out the associated effects of sense and antisense transcripts is difficult. Finally, it is critical to discover to diversify ancestry and sex in genetic association studies to identify more genetic loci underlying MHO. Without experimental follow-up and extensive clinical studies, genotype should not be used as a diagnostic metric. CRISPR editing of alleles in relevant cell types to study *cis*-regulatory effects on genes and phenotypic effects on cells, and work in animal models is necessary to fully annotate these loci. In addition, it is important to identify the indirect and direct effects of discordant variants, as these endocrine tissues are major contributors to peptide and hormone secretion. Further experimental characterization is critical to placing these results in the proper context and providing the basis for personalized interventions for T2D. The predictions at these six loci provide specific hypotheses to be tested, and should they be validated experimentally provide knowledge of the precise mechanisms of uncoupling obesity from T2D risk.

## METHODS

### GWAS-GWAS Colocalization Analysis

GWAS results for T2DadjBMI and WHRadjBMI were obtained from Mahajan et al^66^ and Pulit et al^77^. The set of single nucleotide polymorphisms (SNPs) within 500kb of a genome-wide significant SNP in either GWAS was included in the colocalization test. Rare variants, defined as SNPs reported to have effect allele frequencies of less than 1% in either GWAS, were excluded. Proximal analysis windows (>250kb) were merged, and the colocalization test was performed on these genetic loci with 3 methods: Coloc.abf^27^, Hyprcoloc^78^, and visual inspection of LocusCompare plots^79^.

The default parameters were used for Hyprcoloc. In Coloc.abf, the default parameters for p1 and p2 prior probabilities were used for the individual GWAS hypotheses. The parameter p12, the prior for single variant colocalization, was set to 5e-06 as prescribed by Wallace et. al^27^ to balance false negative and positive results. Loci were considered colocalized if the regional probability of colocalization was greater than 0.70. In Coloc.abf, this was the sum of the PPH3 and PPH4 statistics, and in Hyprcoloc this was the regional probability statistic. Loci that met colocalization criteria in either method were plotted using LocusCompare with the default European ancestry linkage disequilibrium (LD) data from 1000Genomes^80^ and with genome build hg19. This resulted in 121 LocusCompare plots on which visual inspection was performed to verify colocalized genetic association signals. If genetic loci were considered colocalized by at least two of the three colocalization analysis methods, we considered these consensus colocalized loci. We termed this consensus analysis ‘COLOC’.

### Discordant Locus Identification

We obtained the 99% credible set of SNPs from the results of Bayesian Factor Analysis implemented through Coloc.abf at each locus. We calculated the Z-scores for the association test of each genetic variant and the GWAS trait. If the Z-score associated with SNP had the opposite sign for association with WHRadjBMI and T2DadjBMI with respect to the same allele and the p-value for the association with both traits was less than 1e-05, we considered the variant discordant. We then identified in which loci the SNPs were located, and queried haploReg^81^ linkage disequilibrium data with the haploR package in R^82^ to separate signals in the same loci using LD clumping (R^2^>0.50) on the discordant variants.

### Phenome-Wide Association Study

We queried the GWAS meta-analysis associations of glycemic and anthropometric traits for each lead discordant variant in the Type 2 Diabetes Knowledge Portal (T2DKP)^31^. We additionally obtained the summary statistics of abdominal fat MRI scans in the UK Biobank and queried these summary statistics for discordant variants^83^.

### Multi-trait Colocalization Analysis

We obtained GWAS summary statistics for Waist Circumference (WC), Hip Circumference (HC), WHR, WHRadjBMI, T2D, and T2DadjBMI. We extracted summary statistics of variants within genetic loci containing a discordant association signal^66,77^ and performed multi-trait colocalization with Hyprcoloc^78^. We considered an association signal colocalized for multiple traits if Hyprcoloc computed a posterior probability for both body fat distribution traits (WC, HC, WHR, and WHRadjBMI) as well as for T2D or T2DadjBMI.

### Fine-mapping Analysis

We performed variable selection in multiple regression as implemented in the R package SuSiE^84^. This method implements the sum of single-effects models to fine-map the causal variant(s) in a locus. Using the T2DadjBMI and WHRadjBMI GWAS summary statistics and the 1000 Genomes LD data, we performed fine-mapping of loci containing a genetic variant associated with discordant effects on T2DadjBMI and WHRadjBMI. We used the default flag options in SuSiE and performed a sensitivity analysis of the results to a range of priors. We selected causal variants with a PPH4 greater than 0.70.

### GWAS-QTL Colocalization Analysis

We obtained expression quantitative trait locus (eQTL) data from the Genotype-Tissue Expression (GTEx) for 49 tissues^85^, the Stockholm-Tartu Atherosclerosis Reverse Networks Engineering Task (STARNET) cohort for 6 tissues^86^, and the Metabolic Syndrome in Men (METSIM) for subcutaneous adipose tissue^39^. We also obtained subcutaneous adipose tissue splice QTL (sQTL) results from the METSIM cohort. Data sources and further information are detailed in (Figure 2 – source data 2). We extracted the QTL data for each gene or transcript within 1Mb of a discordant locus start or end site and independently colocalized with the T2DadjBMI and WHRadjBMI GWAS using Coloc.abf and Summary-Based Mendelian Randomization (SMR). When implementing Coloc.abf, we considered a signal to be colocalized if PPH4 was greater than 0.50 (a threshold used for GWAS-QTL colocalization in admixed populations^87^). We repeated the analysis in SMR and used a false-discovery rate (FDR) threshold of 5% to control for false positives. We then performed a visual inspection of GWAS-QTL colocalization of plots generated by LocusCompare. If a GWAS-QTL colocalization met these criteria, the proximal gene was termed an effector Gene (eGene).

### fGWAS Annotation Enrichment analysis

We used the functional GWAS (fGWAS)^88^ command-line tool to compute the enrichment of associations in particular genomic and epigenomic regions. We first obtained the chromosome and base-pair position of each variant in the 99% credible set from each of the 79 colocalized loci. We mapped the SNPs to their placement in genomic regions using bed files. We used bed files from tissue-specific chromatin-state data (adipose, liver, pancreatic islet, and skeletal muscle) and genome-level coding region annotations, and mapped SNPs to their presence in these regions. From these maps, we performed enrichment analysis with the complete model of all annotations with the -fine and -xv flags on fGWAS. We used the natural log of the Bayes Factor of the colocalization test and computed the enrichment of SNPs for presence in coding regions to genetic and epigenetic annotations.

### Tissue of Action Analysis

We conducted tissue-of-action (TOA) score analysis using the credible set of SNPs from each of the 79 colocalized loci. TACTICAL computes the TOA score with the SNP-level Bayesian probabilities, the SNP annotation maps, and the annotation enrichment scores. We used the Coloc.abf PPH4 scores for the SNP-level Bayesian probability, the fGWAS annotation enrichment scores, and the SNP annotation maps to compute the tissue of action score at all colocalized loci. We separated independent association signals in the same loci (LD < 0.5) with HaploReg^81^. With TACTICAL^29^, we integrated the credible set of SNPs with the enrichment for genome-level and tissue-specific annotations. We used the default tissue classification thresholds of .20 to classify signals as belonging to a particular TOA and less than .10 difference to classify signals as sharing TOA assignments between multiple tissues.

### Gene Expression and Phenotype Correlation Analysis

For each eGene, we computed the biweight midcorrelation and its significance, as implemented by the Weighted Genetic Coexpression Network Analysis (WGCNA) package^89^, between gene expression with metabolic phenotypes measured in the METSIM cohort^40^. We controlled for false positives with a 5% FDR threshold as implemented by the q-values package in R^90^.

### Adipogenesis Gene Expression Dynamics Analysis

We obtained Simpson-Golabi-Behmel Syndrome (SGBS) preadipocyte adipogenesis time series gene expression data from GEO (accession number GSE76131)^47^. We evaluated the dynamic expression of each adipose tissue eGene by fitting the gene expression over time to a linear model and applying the likelihood ratio test (LRT) to compare the time-dependent models to time-independent null models. We considered an eGene to be dynamically expressed in adipogenesis if the p-value of the LRT was less than 0.05.

## DATA AVAILABILITY

Our analysis pipeline is publicly available on GitHub (https://github.com/aberrations/predicting-functional-mechanisms-discordant-loci). All source data used in our analyses are detailed in source data 1-4.

## SUPPLEMENTARY FIGURE CAPTIONS

**Figure 4-figure supplement 1.**
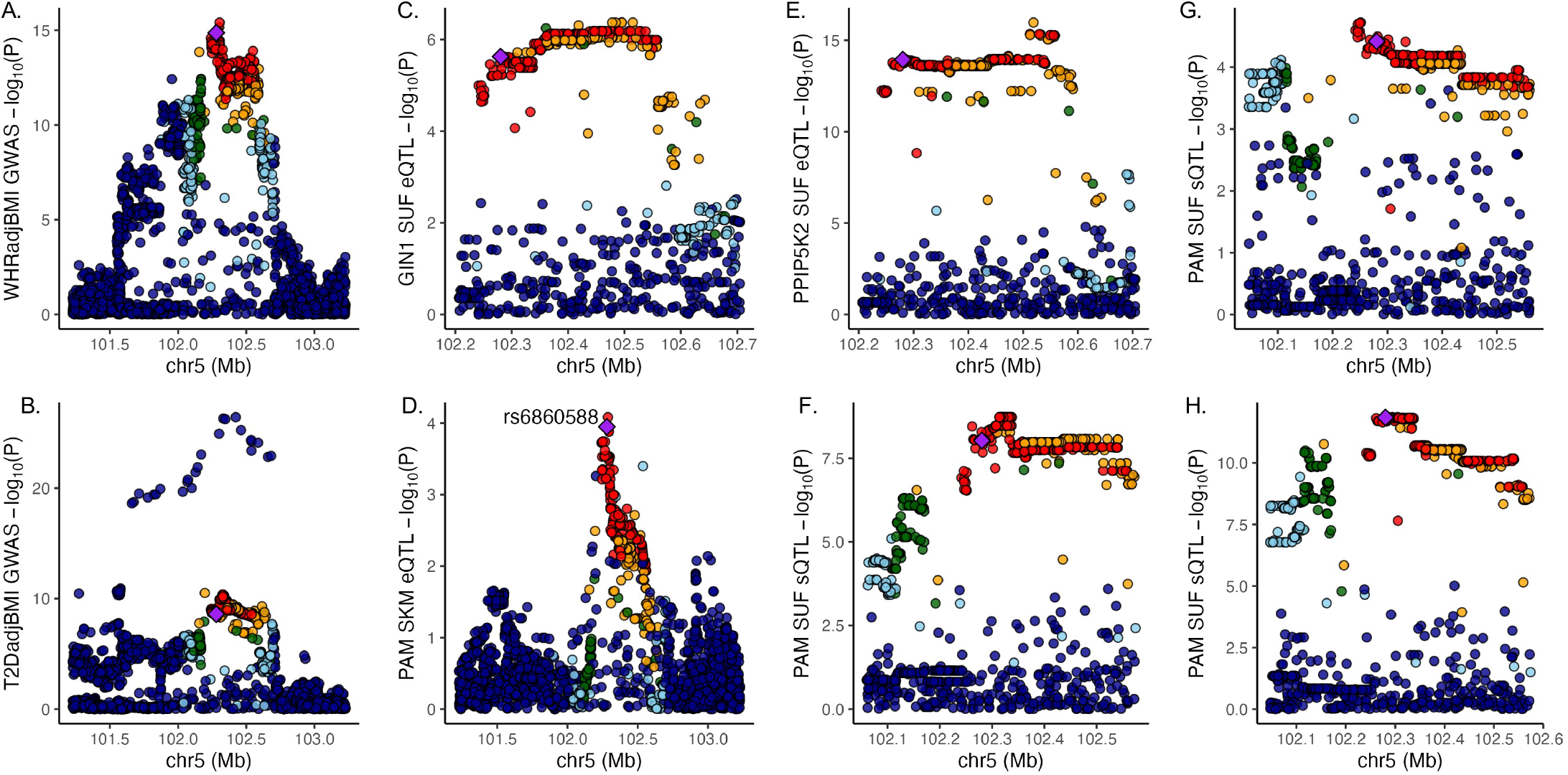
Discordant variant rs6860588 is associated with pleiotropic effects on gene regulation in multiple tissues. Manhattan plot of associations at the 5q21.1 locus containing lead variant rs6860588 for (A) WHRadjBMI GWAS, (B) T2DadjBMI, (C) SAT expression of GIN1, (D) SKM expression of PAM, (E) SAT expression of PPIP5K2 (F) SAT expression of PAM splice variant that skips exon 14, (G) SAT expression of canonical PAM splice junction between exon 13 and 14, and (H) SAT expression of canonical PAM splice junction between exon 14 and 15.

**Figure 4-figure supplement 2.**
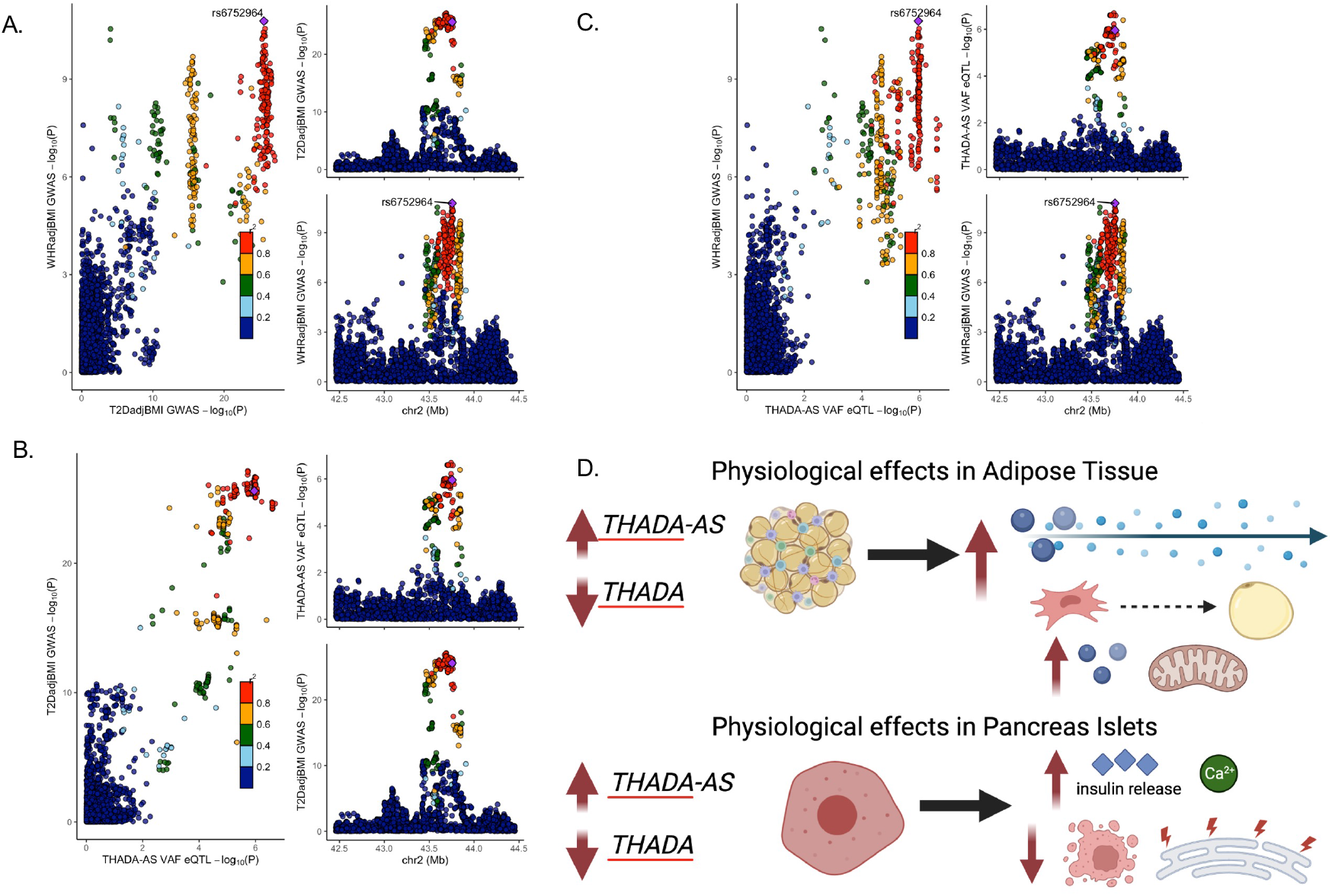
Discordant variant rs6752964 is associated with pleiotropic effects on gene regulation in multiple tissues. LocusCompare plot of associations with rs6752964 for (A) WHRadjBMI GWAS with T2DadjBMI GWAS, (B) T2DadjBMI GWAS with adipose tissue eQTL, and (C) WHRadjBMI GWAS with adipose tissue eQTL. (D) The proposed model reconciles the metabolic discordance observed in 2p21 with the associated lead variant rs6752964. Created with BioRender.com.

## REFERENCES

1. Lusis, A. J. A thematic review series: systems biology approaches to metabolic and cardiovascular disorders. J. Lipid Res. 47, 1887–1890 (2006).

2. McCarthy, M. I. Genomics, Type 2 Diabetes, and Obesity. N. Engl. J. Med. 363, 2339–2350 (2010).

3. Brown, E., Heerspink, H. J. L., Cuthbertson, D. J. & Wilding, J. P. H. SGLT2 inhibitors and GLP-1 receptor agonists: established and emerging indications. The Lancet 398, 262–276 (2021).

4. Blüher, M. Metabolically healthy obesity. Endocr. Rev. 41, 405–420 (2020).

5. Smith, G. I., Mittendorfer, B. & Klein, S. Metabolically healthy obesity: facts and fantasies. J. Clin. Invest. 129, 3978–3989 (2019).

6. Loos, R. J. F. & Kilpeläinen, T. O. Genes that make you fat, but keep you healthy. J. Intern. Med. 284, 450–463 (2018).

7. Klöting, N. et al. Insulin-sensitive obesity. Am. J. Physiol. Endocrinol. Metab. 299, E506–515 (2010).

8. Karelis, A. D. et al. The metabolically healthy but obese individual presents a favorable inflammation profile. J. Clin. Endocrinol. Metab. 90, 4145–4150 (2005).

9. Chen, D. L. et al. Phenotypic Characterization of Insulin-Resistant and Insulin-Sensitive Obesity. J. Clin. Endocrinol. Metab. 100, 4082–4091 (2015).

10. Jennings, C. L. et al. Determinants of insulin-resistant phenotypes in normal-weight and obese Black African women. Obes. Silver Spring Md 16, 1602–1609 (2008).

11. Hayes, L. et al. Do obese but metabolically normal women differ in intra-abdominal fat and physical activity levels from those with the expected metabolic abnormalities? A cross-sectional study. BMC Public Health 10, 723 (2010).

12. Koster, A. et al. Body fat distribution and inflammation among obese older adults with and without metabolic syndrome. Obes. Silver Spring Md 18, 2354–2361 (2010).

13. Emdin, C. A. et al. Genetic association of waist-to-hip ratio with cardiometabolic traits, type 2 diabetes, and coronary heart disease. JAMA - J. Am. Med. Assoc. 317, 626–634 (2017).

14. Gill, D. et al. Risk factors mediating the effect of body mass index and waist-to-hip ratio on cardiovascular outcomes: Mendelian randomization analysis. Int. J. Obes. 45, 1428–1438 (2021).

15. Li, K. et al. Causal associations of waist circumference and waist-to-hip ratio with type II diabetes mellitus: new evidence from Mendelian randomization. Mol. Genet. Genomics MGG 296, 605–613 (2021).

16. Minster, R. L. et al. A thrifty variant in CREBRF strongly influences body mass index in Samoans. Nat. Genet. 48, 1049–1054 (2016).

17. Kim, J.-Y. et al. Obesity-associated improvements in metabolic profile through expansion of adipose tissue. J. Clin. Invest. 117, 2621–2637 (2007).

18. Fathzadeh, M. et al. FAM13A affects body fat distribution and adipocyte function. Nat. Commun. 11, 1465 (2020).

19. Small, K. S. et al. Regulatory variants at KLF14 influence type 2 diabetes risk via a female-specific effect on adipocyte size and body composition. Nat. Genet. 50, 572–580 (2018).

20. Yang, Q. et al. Adipocyte-Specific Modulation of KLF14 Expression in Mice Leads to Sex-Dependent Impacts on Adiposity and Lipid Metabolism. Diabetes 71, 677–693 (2022).

21. Gesta, S. et al. Mesodermal developmental gene Tbx15 impairs adipocyte differentiation and mitochondrial respiration. Proc. Natl. Acad. Sci. U. S. A. 108, 2771–2776 (2011).

22. Loh, N. Y. et al. RSPO3 impacts body fat distribution and regulates adipose cell biology in vitro. Nat. Commun. 11, p(2020).

23. Loh, N. Y. et al. LRP5 regulates human body fat distribution by modulating adipose progenitor biology in a dose- and depot-specific fashion. Cell Metab. 21, 262–273 (2015).

24. Hukku, A. et al. Probabilistic colocalization of genetic variants from complex and molecular traits: promise and limitations. Am. J. Hum. Genet. 108, 25–35 (2021).

25. Wallace, C. Statistical testing of shared genetic control for potentially related traits. Genet. Epidemiol. 37, 802–813 (2013).

26. Wallace, C. et al. Statistical colocalization of monocyte gene expression and genetic risk variants for type 1 diabetes. Hum. Mol. Genet. 21, 2815–2824 (2012).

27. Wallace, C. Eliciting priors and relaxing the single causal variant assumption in colocalisation analyses. PLOS Genet. 16, e1008720 (2020).

28. Hormozdiari, F. et al. Colocalization of GWAS and eQTL Signals Detects Target Genes. Am. J. Hum. Genet. 99, 1245–1260 (2016).

29. Torres, J. M. et al. A Multi-omic Integrative Scheme Characterizes Tissues of Action at Loci Associated with Type 2 Diabetes. Am. J. Hum. Genet. 107, 1011–1028 (2020).

30. Li, Z. et al. Integrating Mouse and Human Genetic Data to Move beyond GWAS and Identify Causal Genes in Cholesterol Metabolism. Cell Metab. 31, 741–754.e5 (2020).

31. Costanzo, M. C. et al. The Type 2 Diabetes Knowledge Portal: An open access genetic resource dedicated to type 2 diabetes and related traits. Cell Metab. 35, 695–710.e6 (2023).

32. McLaren, W. et al. The Ensembl Variant Effect Predictor. Genome Biol. 17, 122 (2016).

33. Civelek, M. & Lusis, A. J. Systems genetics approaches to understand complex traits. Nat. Rev. Genet. 15, 34–48 (2014).

34. Civelek, M. et al. Genetic Regulation of Adipose Gene Expression and Cardio-Metabolic Traits. Am. J. Hum. Genet. 100, 428–443 (2017).

35. Emont, M. P. et al. A single-cell atlas of human and mouse white adipose tissue. Nature 603, 926–933 (2022).

36. Lu, Y. et al. New loci for body fat percentage reveal link between adiposity and cardiometabolic disease risk. Nat. Commun. 7, 10495 (2016).

37. Locke, A. E. et al. Genetic studies of body mass index yield new insights for obesity biology. Nature 518, 197–206 (2015).

38. Hansen, G. T. et al. Genetics of sexually dimorphic adipose distribution in humans. Nat. Genet. 55, 461– 470 (2023).

39. Brotman, S. M. et al. Subcutaneous adipose tissue splice quantitative trait loci reveal differences in isoform usage associated with cardiometabolic traits. Am. J. Hum. Genet. 109, 66–80 (2022).

40. Laakso, M. et al. The Metabolic Syndrome in Men study: a resource for studies of metabolic and cardiovascular diseases. J. Lipid Res. 58, 481–493 (2017).

41. Volarevic, V., Al-Qahtani, A., Arsenijevic, N., Pajovic, S. & Lukic, M. L. Interleukin-1 receptor antagonist (IL-1Ra) and IL-1Ra producing mesenchymal stem cells as modulators of diabetogenesis. Autoimmunity 43, 255–263 (2010).

42. Luotola, K. IL-1 Receptor Antagonist (IL-1Ra) Levels and Management of Metabolic Disorders. Nutrients 14, 3422 (2022).

43. Grossmann, V. et al. Profile of the Immune and Inflammatory Response in Individuals With Prediabetes and Type 2 Diabetes. Diabetes Care 38, 1356–1364 (2015).

44. Herwald, H. & Egesten, A. C-Reactive Protein: More than a Biomarker. J. Innate Immun. 13, 257–258 (2021).

45. Li, S., Liberman, L. M., Mukherjee, N., Benfey, P. N. & Ohler, U. Integrated detection of natural antisense transcripts using strand-specific RNA sequencing data. Genome Res. 23, 1730 (2013).

46. Achari, A. E. & Jain, S. K. Adiponectin, a Therapeutic Target for Obesity, Diabetes, and Endothelial Dysfunction. Int. J. Mol. Sci. 18, 1321 (2017).

47. Nassiri, I. et al. Systems view of adipogenesis via novel omics-driven and tissue-specific activity scoring of network functional modules. Sci. Rep. 6, 28851 (2016).

48. Anderson, W. D. et al. Sex differences in human adipose tissue gene expression and genetic regulation involve adipogenesis. Genome Res. 30, 1379–1392 (2020).

49. Li, W., Wang, L., Wu, Y., Yuan, Z. & Zhou, J. Weighted gene co-expression network analysis to identify key modules and hub genes associated with atrial fibrillation. Int. J. Mol. Med. 45, 401–416 (2020).

50. Thomsen, S. K. et al. Type 2 diabetes risk alleles in PAM impact insulin release from human pancreatic β-cells. Nat. Genet. 50, 1122–1131 (2018).

51. Merkler, D. J. C-terminal amidated peptides: production by the in vitro enzymatic amidation of glycine-extended peptides and the importance of the amide to bioactivity. Enzyme Microb. Technol. 16, 450–456 (1994).

52. Chen, Y.-C. et al. PAM haploinsufficiency does not accelerate the development of diet- and human IAPP-induced diabetes in mice. Diabetologia 63, 561–576 (2020).

53. Zieliński, M. et al. Expression of recombinant human bifunctional peptidylglycine α-amidating monooxygenase in CHO cells and its use for insulin analogue modification. Protein Expr. Purif. 119, 102– 109 (2016).

54. Gu, C. et al. Metabolic supervision by PPIP5K, an inositol pyrophosphate kinase/phosphatase, controls proliferation of the HCT116 tumor cell line. Proc. Natl. Acad. Sci. U. S. A. 118, e2020187118 (2021).

55. Gu, C. et al. KO of 5-InsP7 kinase activity transforms the HCT116 colon cancer cell line into a hypermetabolic, growth-inhibited phenotype. Proc. Natl. Acad. Sci. 114, 11968–11973 (2017).

56. Badodi, S. et al. Inositol treatment inhibits medulloblastoma through suppression of epigenetic-driven metabolic adaptation. Nat. Commun. 12, 2148 (2021).

57. Caleyachetty, R. et al. Metabolically Healthy Obese and Incident Cardiovascular Disease Events Among 3.5 Million Men and Women. J. Am. Coll. Cardiol. 70, 1429–1437 (2017).

58. Rey-López, J. P., de Rezende, L. F., de Sá, T. H. & Stamatakis, E. Is the metabolically healthy obesity phenotype an irrelevant artifact for public health? Am. J. Epidemiol. 182, 737–741 (2015).

59. Blüher, M. Obesity: The myth of innocent obesity. Nat. Rev. Endocrinol. 13, 691–692 (2017).

60. Kilpeläinen, T. O. et al. Genetic variation near IRS1 associates with reduced adiposity and an impaired metabolic profile. Nat. Genet. 43, 753–760 (2011).

61. Lotta, L. A. et al. Integrative genomic analysis implicates limited peripheral adipose storage capacity in the pathogenesis of human insulin resistance. Nat. Genet. 49, 17–26 (2017).

62. Censin, J. C. et al. Causal relationships between obesity and the leading causes of death in women and men. PLOS Genet. 15, e1008405 (2019).

63. Dale, C. E. et al. Causal Associations of Adiposity and Body Fat Distribution With Coronary Heart Disease, Stroke Subtypes, and Type 2 Diabetes Mellitus: A Mendelian Randomization Analysis. Circulation 135, 2373–2388 (2017).

64. Klonoff, D. C. Personalized Medicine for Diabetes. J. Diabetes Sci. Technol. Online 2, 335–341 (2008).

65. Williams, D. M., Jones, H. & Stephens, J. W. Personalized Type 2 Diabetes Management: An Update on Recent Advances and Recommendations. Diabetes Metab. Syndr. Obes. Targets Ther. 15, 281–295 (2022).

66. Mahajan, A. et al. Fine-mapping type 2 diabetes loci to single-variant resolution using high-density imputation and islet-specific epigenome maps. Nat. Genet. 50, 1505–1513 (2018).

67. Zeggini, E. et al. Meta-analysis of genome-wide association data and large-scale replication identifies additional susceptibility loci for type 2 diabetes. Nat. Genet. 40, 638–645 (2008).

68. Grarup, N. et al. Association testing of novel type 2 diabetes risk alleles in the JAZF1, CDC123/CAMK1D, TSPAN8, THADA, ADAMTS9, and NOTCH2 loci with insulin release, insulin sensitivity, and obesity in a population-based sample of 4,516 glucose-tolerant middle-aged Danes. Diabetes 57, 2534–2540 (2008).

69. Moraru, A. et al. THADA Regulates the Organismal Balance between Energy Storage and Heat Production. Dev. Cell 41, 72–81.e6 (2017).

70. Zhang, Y. et al. THADA inhibition in mice protects against type 2 diabetes mellitus by improving pancreatic β-cell function and preserving β-cell mass. Nat. Commun. 14, 1020 (2023).

71. Simonis-Bik, A. M. et al. Gene variants in the novel type 2 diabetes loci CDC123/CAMK1D, THADA, ADAMTS9, BCL11A, and MTNR1B affect different aspects of pancreatic β-cell function. Diabetes 59, 293– 301 (2010).

72. Brantl, S. Antisense-RNA regulation and RNA interference. Biochim. Biophys. Acta 1575, 15–25 (2002).

73. Faghihi, M. A. & Wahlestedt, C. Regulatory roles of natural antisense transcripts. Nat. Rev. Mol. Cell Biol. 10, 637–643 (2009).

74. Wight, M. & Werner, A. The functions of natural antisense transcripts. Essays Biochem. 54, 91–101 (2013).

75. Chen, Z. et al. Functional Screening of Candidate Causal Genes for Insulin Resistance in Human Preadipocytes and Adipocytes. Circ. Res. 126, 330–346 (2020).

76. Lin, Q.-Z. et al. Sex-specific association of the peptidase D gene rs731839 polymorphism and serum lipid levels in the Mulao and Han populations. Int. J. Clin. Exp. Pathol. 7, 4156–4172 (2014).

77. Pulit, S. L. et al. Meta-Analysis of genome-wide association studies for body fat distribution in 694 649 individuals of European ancestry. Hum. Mol. Genet. 28, 166–174 (2019).

78. Foley, C. N. et al. A fast and efficient colocalization algorithm for identifying shared genetic risk factors across multiple traits. Nat. Commun. 12, 592238 (2021).

79. Liu, B., Gloudemans, MichaelJ., Rao, A. S., Ingelsson, E. & Montgomery, S. B. Abundant associations with gene expression complicate GWAS follow-up. Nat. Genet. 51, 768–769 (2019).

80. Fairley, S., Lowy-Gallego, E., Perry, E. & Flicek, P. The International Genome Sample Resource (IGSR) collection of open human genomic variation resources. Nucleic Acids Res. 48, D941–D947 (2020).

81. Ward, L. D. & Kellis, M. HaploReg v4: systematic mining of putative causal variants, cell types, regulators and target genes for human complex traits and disease. Nucleic Acids Res. 44, D877–881 (2016).

82. Zhbannikov, I. Y., Arbeev, K., Ukraintseva, S. & Yashin, A. I. haploR: an R package for querying webbased annotation tools. F1000Research 6, 97 (2017).

83. Liu, Y. et al. Genetic architecture of 11 organ traits derived from abdominal MRI using deep learning. eLife 10, e65554.

84. Wang, G., Sarkar, A., Carbonetto, P. & Stephens, M. A simple new approach to variable selection in regression, with application to genetic fine mapping. J. R. Stat. Soc. Ser. B Stat. Methodol. 82, 1273–1300 (2020).

85. Aguet, F. et al. Genetic effects on gene expression across human tissues. Nature 550, 204–213 (2017).

86. Franzén, O. et al. Cardiometabolic risk loci share downstream cis- and trans-gene regulation across tissues and diseases. Science 353, 827–830 (2016).

87. Gay, N. R. et al. Impact of admixture and ancestry on eQTL analysis and GWAS colocalization in GTEx. Genome Biol. 21, 233 (2020).

88. Pickrell, J. K. Joint analysis of functional genomic data and genome-wide association studies of 18 human traits. Am. J. Hum. Genet. 94, 559–573 (2014).

89. Langfelder, P. & Horvath, S. WGCNA: an R package for weighted correlation network analysis. BMC Bioinformatics 9, 559 (2008).

90. A direct approach to false discovery rates - Storey - 2002 - Journal of the Royal Statistical Society: Series B (Statistical Methodology) - Wiley Online Library. https://rss.onlinelibrary.wiley.com/doi/full/10.1111/1467-9868.00346.

